# Phosphorylation of interfacial phosphosite leads to increased binding of Rap-Raf complex

**DOI:** 10.1101/2021.12.19.473331

**Authors:** T Devanand, Susmita Ghosh, Prasanna Venkatraman, Satyavani Vemparala

## Abstract

The effect of phosphorylation of a serine residue in the Rap protein, residing at the complex interface of Rap-Raf complex is studied using atomistic molecular dynamics simulations. As the phosphosite of interest (SER39) is buried at the interface of the Rap-Raf complex, phosphorylation of only Rap protein was simulated and then complexed with the RBD of Raf for further analysis of complex stability. Our simulations reveal that the phosophorylation increases the binding of complex through strong electrostatic interactions and changes the charge distribution of the interface significantly. This is manifested as an increase in stable salt-bridge interactions between the Rap and Raf of the complex. Network analysis clearly shows that the phosphorylation of SER39 reorganizes the community network to include the entire region of Raf chain, including, Raf L4 loop potentially affecting downstream signalling.

## I. INTRODUCTION

Rap proteins belong to the small Ras-like GTPase family which are involved in many crucial cellular functions such as cell adhesion, cell-cell junction formation and regulation of the actin cytoskeleton^1–4^. These GTPases act like molecular switches which are active if GTP ligand is bound to the protein and inactive when GTP is replaced with GDP. The rates of activity of these molecular switches are determined by factors like guanine exchange factors(GEFs), which catalyze Rap/Ras proteins to GTP-bound active state and GTPase activating proteins(GAPs) which enhance the intrinsic hydrolysis of bound GTP to GDP.^5^ Ras-like GTPase proteins regulate important signalling pathways such as mitogen activated protein kinase (MAPK) MAPK/ERK and PI3K^6^ and such regulation is determined by the effector proteins with which they interact and Raf-1 is one of the most important effector molecules for Ras family of proteins^7^. Raf is the first kinase in the MAPK cascade and interacts with Ras-like GTPases via the Ras binding domain (RBD). Ras-like GTPases and their mutations have been identified as some of the key drivers of oncogenesis in several human tumors and intense efforts have been made to understand the basis of modulation of their activation and signaling via different methods.

Direct inhibition of the Ras-like GTPase via small molecules has not been very successful and these proteins have been termed as notoriously undruggable, in terms of inability to identify a well-defined binding pocket for drug molecules^8,9^. Many recent efforts have been focusing on alternative ways to affect the function and activity of Ras-family of proteins including inhibitors of Ras downstream effector signalling, Ras-membrane interaction blockers and small molecules that disrupt Ras-Raf interactions^8,10–12^. A promising direction to explore is to target the many post translational modification (PTM) sites^13,14^ on either Ras or other competing proteins like Rap and indirectly affect the Ras-Raf interaction. Rap1 proteins^15–21^ are identified as a suppressors of Ras activity in screening assays, due to their ability to competitively bind (in the presence of GTP ligand) to downstream Raf without activating it and inhibiting the Ras-Raf interaction hence disrupting the signal transmission along the MAPK pathway^17,22–26^.

Reversible PTMs like phosphorylation^27^ are integral part of many signaling pathways and it is no surprise that ligand-based activity switching proteins like Ras-like GTPases have been shown to possess multiple PTM sites, which can be potentially targeted for the development of small molecule inhibitors that might alter the function of these GTPases. This alternate paradigm of limiting the activity of Ras-like GTPases becomes especially relevant given the failure to find successful direct inhibitors of Ras proteins. We have recently showed that the phosphorylation of a single serine residue (SER11), located proximally to the bound GTP ligand triggers a cascade of interactions resulting in enhanced binding affinity of the Rap1-Raf complex.^28^. A possible implication of this increased binding affinity of the complex can potentially inhibit Ras-Raf interaction by making Raf less available to Ras proteins thus affecting the downstream signalling pathway. Our simulations also demonstrated that a new pocket is revealed in the phosphorylated Rap1 protein and a possible allosteric pathway opens up, spanning more than 40°A between the phosphorylated SER11 and the L4 loop of RBD of Raf. Experimental studies^29^ revealed more possible phosphosites in Rap1 protein including SER39 located on the Switch I loop, which is implicated in many complexes of Rap1 and its corresponding effector proteins^30,31^. Switch I loop plays an important role in many interactions of Rap1 with the Ras binding domain(RBD) regions of effector or regulator protein like Raf and the inactive(GDP-bound) to active(GTP-bound) switching of protein is accompanied largely by conformational changes in Switch I and Switch II loops. In addition, a large number of phosphoproteomics studies suggest that a significant number of phosphorylations occur on phosophosites which are buried^32–36^. The phosphosites could be buried within a protein or at the complex interface and it is intriguing what dynamical aspects of protein conformation are involved in exposing such sites to solvent/kinase of interest. In particular, SER39 has been shown to contribute significantly the kinetic differences of interactions of an adapter protein RAPL with RAP1 and RAP2 proteins, further suggesting its key role at complex interface^37^.

In this work, we explore the effects of phosphorylation of SER39 residue which is located on functionally relevant Switch I loop and at the Rap-Raf complex interface. The buried location of this phosphosite at the interface of the complex presents two issues to be probed:(1) what are the effects of phosphorylation of a residue at the complex interface on the structure of both Rap and Rafã (2) how does the phosphorylation affect the interactions at the complex interface and affect the binding of Rap-Raf complexã

## II. MODEL AND METHODS

### A. MD Simulations

To understand the role of phosphorylation of SER39 on the dynamics of Rap-Raf complex, we performed three simulations: (1) a control simulation in which no residue in the complex was phosphorylated (referred to as CTRL henceforth) (for 400 ns), (2) only the Rap protein with phosphorylated SER39 was simulated for 300 ns and the RBD-Raf was later docked on to this phosphorylated Rap and simulated for 550 ns (referred to as PSER39 henceforth). The initial structure for all the simulations is taken from Protein Data Bank with id 1C1Y^31^, with a resolution of 2.2°A, and contains Rap1A (referred to as Rap henceforth) liganded with a GTP-analogue molecule, Mg^2+^ ion and RBD (Ras binding Domain) region of Raf protein variant c-Raf1 (referred to as Raf henceforth). The missing atoms were modelled using MODELLER software^38^ and the appropriate protonation states for all ionizable residues were determined using PDB2PQR server^39^. The GTP analogue in the crystal structure, GppNHp was replaced by GTP molecule for the simulations. All atom molecular dynamics (MD) simulations were performed on both the systems using NAMD^40^ software and CHARMM36^41,42^ forcefield. VMD software^43^ was used to visualize the system, setup the basic systems and to perform data analysis etc. Each system was solvated in a water box (using TIP3P^44^ water model) and overall charge neutrality was achieved through the addition of appropriate counter ions and to maintain 150 mM salt concentration. The systems were then subjected to energy minimization runs using the conjugate gradient method for 5000 steps, followed by MD simulation runs in NPT ensemble with the integrator time step set to 2fs during equilibration and production runs. The NoséHoover-Langevin piston with a decay period of 100fs and a damping time of 50fs was used to maintain a constant pressure of 1atm.^45,46^ Berendsen thermostat^47^ was used to control temperature at 298 K. A cut-off distance of 12°A was used to compute all short-range van der Waals (VDW) interactions and the long-range electrostatics interactions was treated with the Particle Mesh Ewald(PME) method^48,49^. Electrostatic potential maps of protein structures of CTRL and PSER39 systems constructed using PDB2PQR/APBS(Adaptive Poisson Boltzman Solver) server^39,50,51^. and visualized using Chimera/PyMOL software. MM-GBSA(Molecular Mechanics-Generalised Born Surface Area) calculations were done to estimate the binding free energy of Rap-Raf complex with and without phosphorylation of SER39 residue. We followed the same methodology as described in^28^.

*NetworkView* ^52,53^ plugin in VMD is used to perform the community network analysis on the MD trajectory data. Each C_*α*_ atom of protein residues is represented as a node in this network analysis, with dynamically evolving contacts as possible edges. A contact is defined if any two non-bonded protein residues are within 4.5°*A* for atleast 75% of the trajectory considered (last 50ns MD simulation trajectory CTRL and PSER39 systems). The software gncommunities^53^ was used and the community detection analysis utilizes Girvan-Newman algorithm^54^.

## III. RESULTS

### A. Effect of phosphorylation of SER39 on complex structure

In this section, we probe the effect of phosphorylation of the complex interface residue SER 39 on the overall structure of the complex. As can be seen in Fig. 1, the complex interface is populated with polar and charged residues (both positive and negative). Phosphorylation of SER39 (introduction of negative charges in such an environment) is expected to result in local rearrangement of charged residue and possible the loops containing such residues. To quantify such rearrangement, as compared to the crystal structure, the root mean squared deviation (RMSD) of each residue (averaged over last 50 ns of simulation) is plotted in Fig. 2 for both unphosphorylated (CTRL) and phosphorylated (PSER39) systems.

**FIG. 1.**
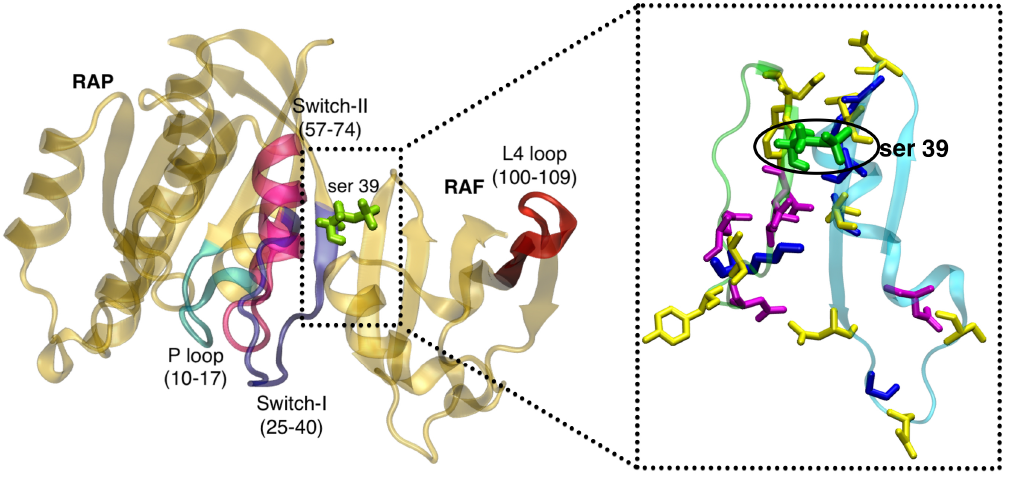
Rap-Raf protein complex (crystal structure, PDB ID 1C1Y) showing the location of important functional loops like P-loop, Switch I, Switch II and RBD loop regions and the interfacial phosphosite SER39. The inset shows the blowup of the Rap-Raf interface and positioning of polar(yellow), positively charged (blue) and negatively charged (purple) interfacial residues.

**FIG. 2.**
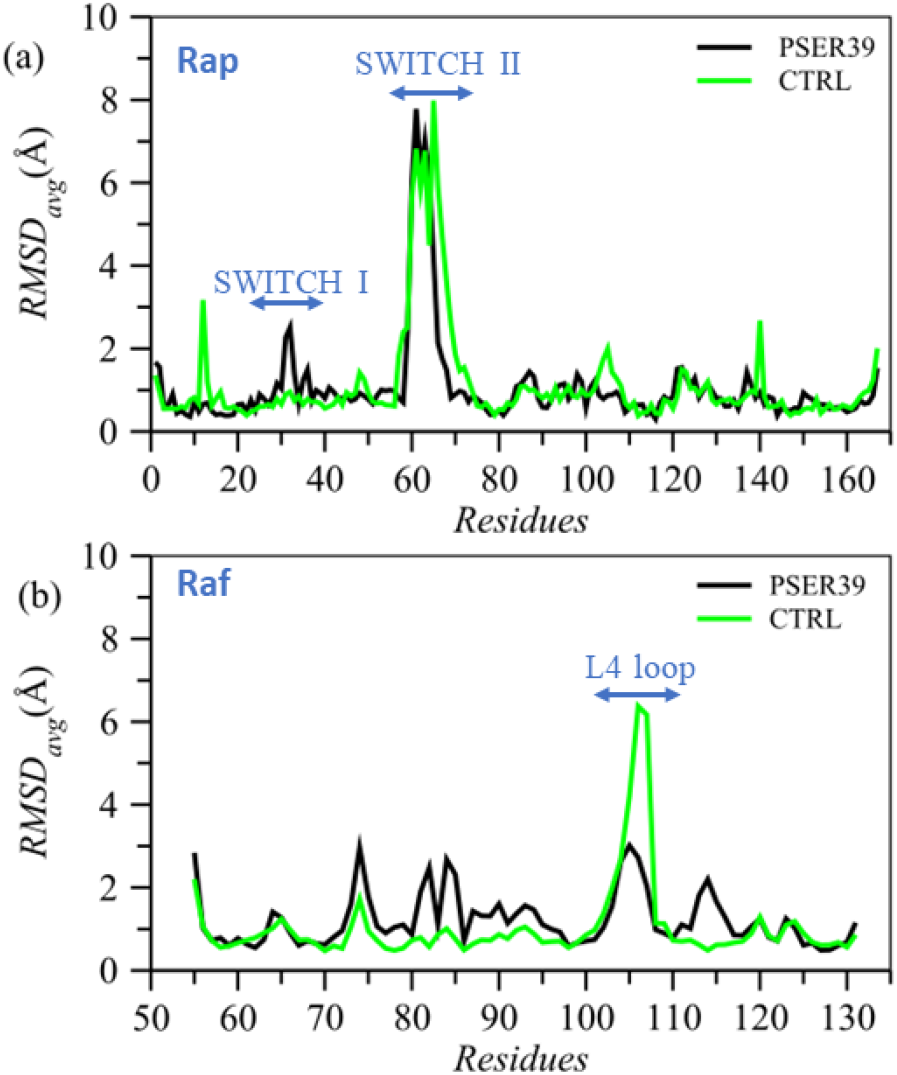
Average of Root mean squared deviations of the residues of the complex with and without phosphorylation of the interface SER39. The values are averaged over of last 50 ns of simulation data.

Given the location of SER39 on Rap protein (surface residue), the phosphorylation does not alter the conformation of most of the Rap protein except for the Switch I loop, where SER39 is located. However, many residues on Raf protein undergo change in conformation as can be seen from overall change in the RMSD values for Raf protein. In particular loops containing residues 60 to 100 (particularly ARG67, ARG89, LYS84, ASP59) and the functionally relevant L4 loop, which is more than 30°*A* from the complex interface. Looking at the mobility of functional loops, via root mean square fluctuations (RMSF) in Supplementary figure 1, it can be seen that the phosphorylation of SER39 has no significant effect on the dynamics of loops like Switch I, Switch II in PSER39 system. The phosphorylation of SER39 significantly increases the accessibility of the residue to water molecules, which is manifested in significant increase in the solvent accessible surface area (SASA) of the pSER39 residue (Figure 3). Earlier studies^55,56^ have shown that SASA of many bound proteins is higher than their unbound counterparts, suggesting that the stability of protein-protein complex can be related to increase in the SASA values. The change in SASA values has also been shown to be an indicator of conformational change^57^, and the observed increase of SASA of SER39 is inline with the results of conformational change in Fig. 2. In Fig. 3, the interface residue conformations and the number of water molecules within 4°*A* of SER39 residue are shown. It can be seen that in the case of PSER39 system, the positively charged amino acids such as ARG41, ARG67 undergo significant conformational changes to face at the complex interface, most likely to counter the negative charges introduced when SER39 residue is phosphorylated. There are also significantly higher number of bridging water molecules between the positively charged and negatively charged residues at the complex interface. To visualize the effect of these conformational changes at the interface, the electrostatic potential map of the Rap-Raf complex using the last frame of both CTRL and PSER39 systems is shown in Fig.4. The electrostatic potential maps show dramatic differences, particularly at the complex interface and in Raf protein, as a response to the introduction of negative charges at the complex interface via phosphorylation. The phosphorylated SER39 residue itself undergoes a conformational change and to assess the conformational change of PSER39, the backbone dihedral angle was measured and it changes from 94 ± 0.14 to 74 ± 0.12 (Data shown in Supplementary figure 2).

**FIG. 3.**
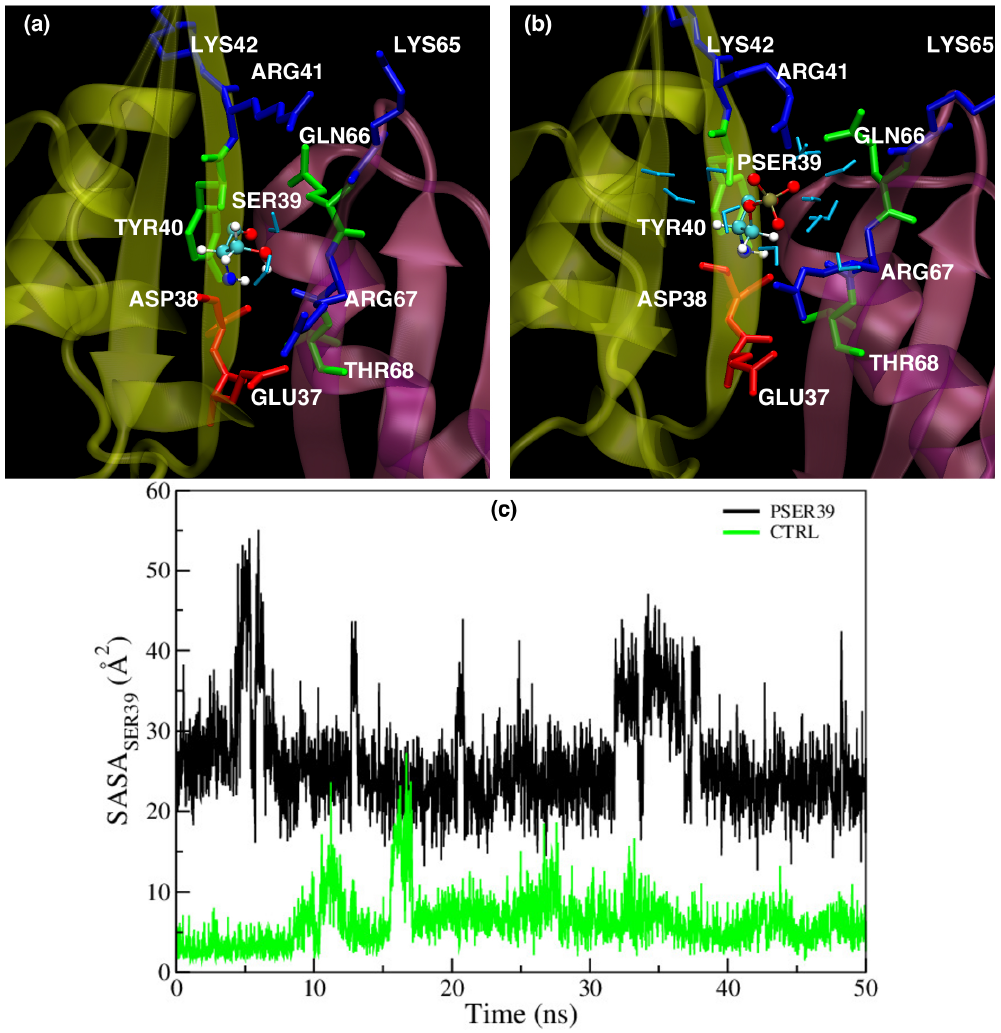
Hydration of Rap-Raf complex interface: complex interface showing relevant residues and water molecules (stick, cyan) within 4*Å* of SER39 residue in (a) CTRL and (b)PSER39 systems. (c) Solvent accessible surface area of SER39 with and without phosphorylation in last 50 ns of simulations.

**FIG. 4.**
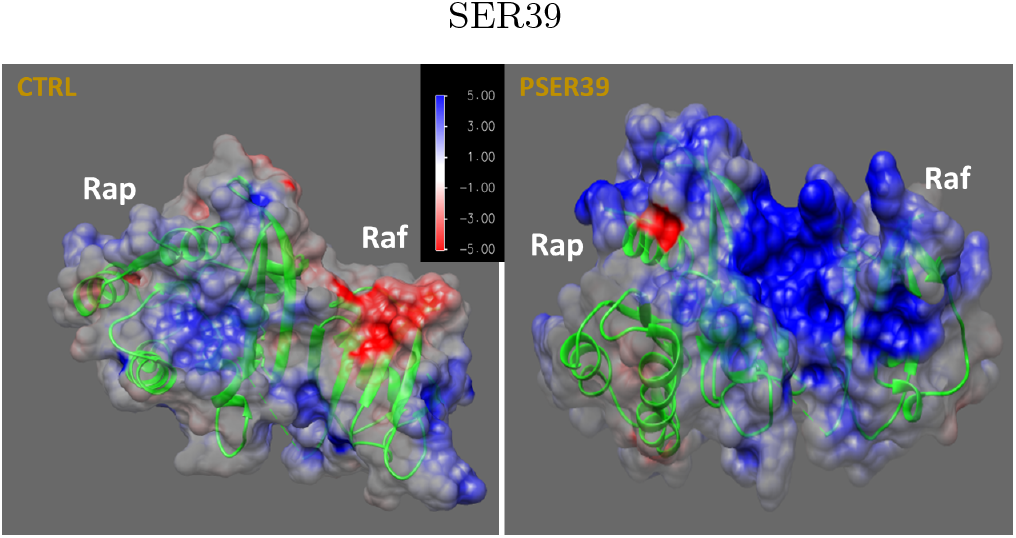
Electrostatic potential map of the complex with and without phosphorylation of the interface serine,

### B. Interactions at the complex interface

As shown in Fig1, the Rap-Raf complex interface is dominated by many polar and charged residues. Herein we explore the emergence of electrostatic interactions between Rap and Raf by monitoring the salt bridge and hydrogen bond interactions across the complex interface. The distribution of hydrogen bonds between Rap and Raf are measured over the last 100ns of simulation for both CTRL and PSER39 systems. The data (Supplementary figure 3) shows an increase in average number of hydrogen bonds between the two proteins when interfacial SER39 is phosphorylated. Previous studies have shown that propensity of salt bridge formation and hydrogen bonding at the interface is significantly higher and can contribute to protein complex stability^58,59^. In Fig.5, time evolution of three possible salt bridges between Rap and Raf protein for CTRL and PSER39 for last 100 ns is shown. For PSER39 system, salt bridge interactions remain stable suggesting increased electrostatic attraction at the complex interface, compared to the CTRL system, where no stable salt bridge interactions exist. The phosphorylated SER39 forms a stable salt bridge with ARG67 of. Raf protein, which is absent in the unphosphorylated. complex. The other two residues ASP33 and GLU37 are also present on the Switch I loop and emergence of stable salt bridge interactions with residues on Raf can influence the conformation, as seen in Fig. 2. In Fig 5(d,e), the conformations of three interfacial residues SER39, ARG67 and GLU37 are shown for both phosphorylated and unphosphorylated systems showing the significant change in the conformations of the interfacial residues when SER39 is phosphorylated.

**FIG. 5.**
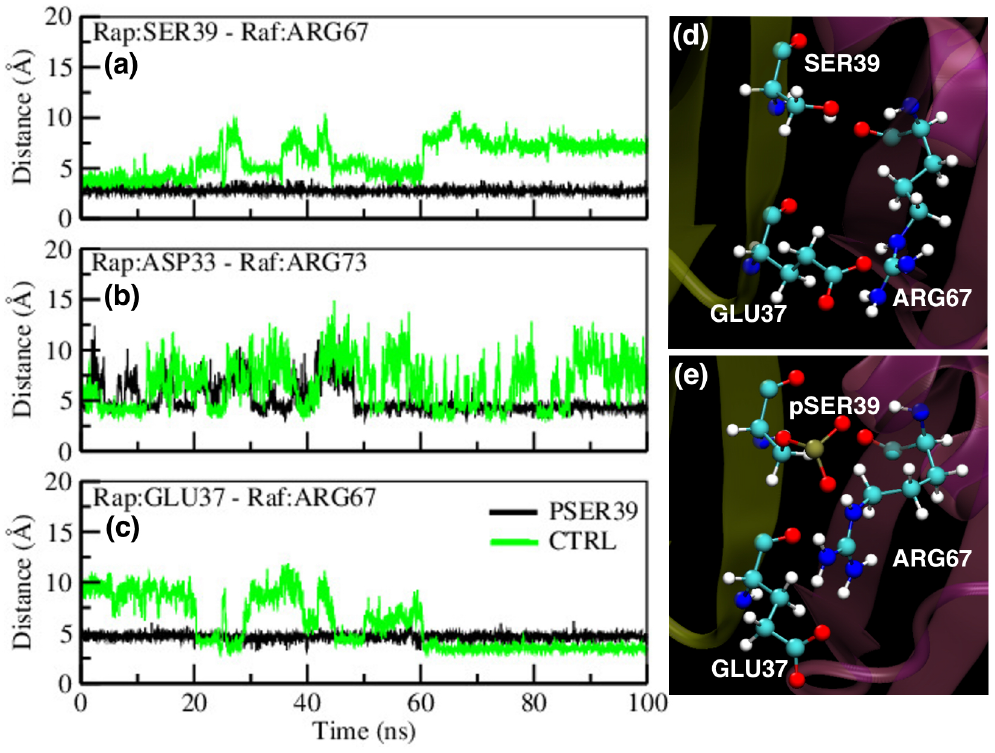
Evolution of Interfacial interaction between Rap and Raf with phosphorylation of SER39 in the form of stable and long-lived salt bridge between pSER39(Rap) and ARG67(Raf). The same interaction for the control case is shown in green.

The effect of phosphorylation on the binding free energy of the Rap-Raf complex is calculated using MMGBSA technique, as described in the Methods section and the results are shown in Table 1. The calculations are performed over last 100ns of both CTRL and PSER39 systems. The binding energy values clearly suggest that the complex has more attractive binding energy when the SER39 residue is phosphorylated indicating an increased binding of Rap-Raf complex in line with our previous results. The electrostatic term contributes the most to this increase and is consistent with increased interactions at the interface as seen in formation of long-surviving electrostatic interactions between phosphorylated SER39 residue of Rap with ARG67 residue of Raf protein shown previously. We have also computed the non-bonded interactions energies between Rap and Raf using the MD trajectories (Supplementary Figure 4), which shows a consistent increase in the attractive electrostatic interactions when the SER39 is phosphorylated. To further understand the global effects of such increased binding of the complex with phosphorylation of SER39, community network analysis was performed. Community network analysis^60^ is a tool to identify possible signaling or interacting pathways between residues of protein, based on molecular dynamics trajectories.^53,54,61–63^. The results of community network analysis are shown in Fig 6 and indicates the emergence of a single dominant community spanning the complex interface and Raf protein in the case of PSER39 system. This implies that the changes in the interface can be transmitted allosterically throughout the Raf protein, possibly affecting the downstream signalling and consistent with the increased binding energy of the complex. The phosphorylation of the interfacial SER39 residue thus has significant effect on increasing the binding of the complex and also possibly trap the Raf protein competitively, making it less available to other proteins such as Ras.

**TABLE I.** Free energy contribution(last 100ns) of GTP liganded simulations.

| Contribution<br>(KCal/mol) | CTRL<br>300-400ns | GTP-PSER39<br>450-550ns |
| --- | --- | --- |
| $\Delta G_{bind}$ | $-62.88 \pm 8.59$ | $-67.52 \pm 7.47$ |
| $\Delta E_{elec}$ | $-498.66 \pm 109.20$ | $-638.40 \pm 48.94$ |
| $\Delta E_{vdW}$ | $-45.88 \pm 5.69$ | $-31.13 \pm 5.61$ |
| $\Delta G_{solv}$ | $481.66 \pm 104.23$ | $602.01 \pm 45.01$ |
| $\Delta G_{solv-np}$ | $-8.95 \pm 0.54$ | $-9.24 \pm 0.47$ |
| $\Delta G_{solv-polar}$ | $490.62 \pm 104.17$ | $611.25 \pm 45.00$ |

**FIG. 6.**
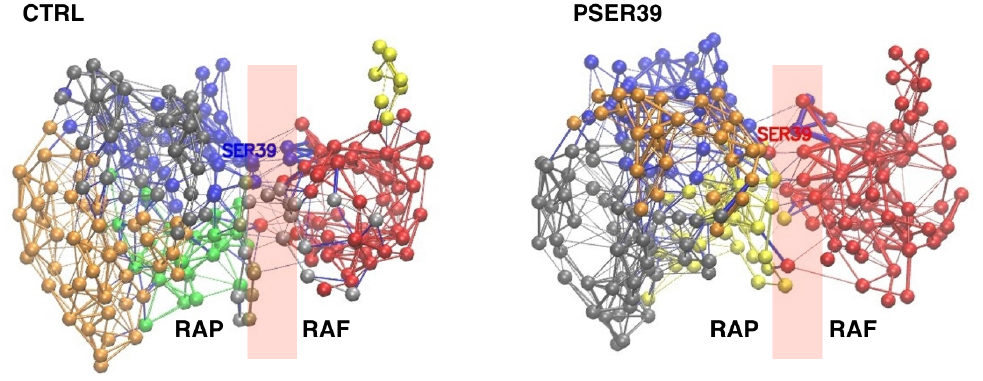
Optimal community network obtained from the correlation map.

## IV. DISCUSSION AND CONCLUSION

Post translational modifications, particularly reversible phosphorylation, can be effectively used to influence many protein-protein interactions and affect signalling pathways in which these proteins participate^64,65^. Many phosphoproteomic studies have suggested that a majority of proteins have phosphosites which can be reversibly phosphorylated by kinases and phosphotases^32,34,66,67^. Studies have also revealed that many such phosphosites are located at the proteinprotein complex interfaces and can affect the binding and complex stability on phosphorylation^68–74^. As can be envisaged, phosphorylation (and subsequent introduction of negative charges) at such interfaces can have significant effects on the conformation, electrostatic interactions and consequently modulate binding and stability of protein-protein complexes.

One of the key goals is to understand how to address the issue of non-druggability of oncoproteins like Ras. A possible direction in which efforts are being made is to address this problem in a unique way: how to reduce the binding of Ras and effector proteins like Raf, which can mitigate the effects of oncogenic forms of Ras in participating the crucial MAPK signalling pathways. In our previous work^28^, we demonstrated how phosphorylation of another predicted phosphosite SER11, which is proximally located near the bound nucleotide, but not at the complex interface, can affect the dynamics of the Switch-I and Switch-II loops via cascading interaction network changes. That study also demonstrated possible longrange allosteric effects of phosphorylation, seen in other systems as well^71,75–77^. The study also showed that the phosphorylation of the SER11 residue increases the binding of the Rap-Raf complex, suggesting that this could be an avenue to make Raf less available to Ras, using the phosphorylation of the antagonist Rap.

Herein we examine the effect of phosphorylation of an interfacial phosphosite SER39, located on functionally relevant Switch-I, in Rap protein on the binding of Raf kinase (RBD) and the consequent complex stability. Our results show that the phosphorylation of SER39 increases the binding of the complex. This is achieved via increased electrostatic attractive interactions between Rap and Raf via salt bridges and hydrogen bond interactions. The introduction of negative charges at the interface, via phosphorylation, results in local conformational changes of charged residues and altering the electrostatic network at the interface. This is also manifested as significant changes in the electrostatic potential maps. The binding energy calculations and the coarse grained community network calculations confirm the increases interactions between the two proteins in the complex. The increased binding energy via phosphorylation of SER39, here, is similar to the increased binding energy when another (non-interfacial) phosphosite SER11 is phosphorylated^28^. Interestingly both SER11 and SER39 are buried phosphosites and it is expected that the inherent protein fluctuations can potentially expose these sites making them kinase accessible for phosphorylation.The results here clearly demonstrate that targeting the protein-protein interface interactions via PTMs like phosphorylation can be an effective tool in increasing the binding of a complex. This can be a potential tool in trapping certain effector proteins by antagonists, making them less available to their primary targets, disrupting signalling pathways, especially in GTPases like Ras, which are notoriously undruggable.

## Supporting information

Supplemetal Figure 1, 2, 3, 4 and will be used for the link on the preprint site

