## Supplementary material for "Phosphorylation of interfacial phosphosite leads to increased binding of Rap-Raf complex": Supplemetal Figure 1, 2, 3, 4 and will be used for the link on the preprint site

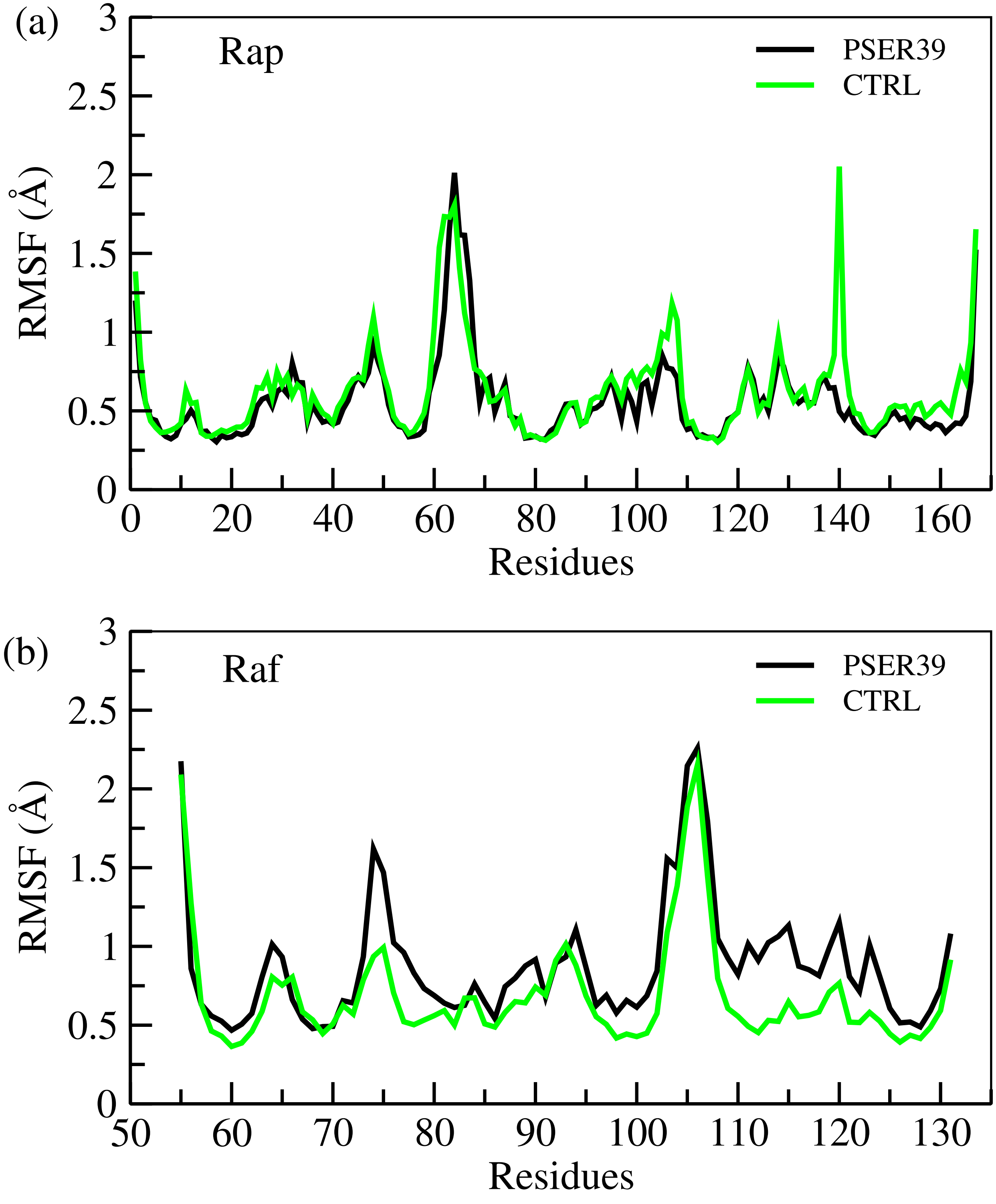


FIG. 1. The RMSF of residues of (a)Rap and (b)Raf $C_\alpha$ atoms averaged over last 100 ns of simulation for PSER39 and CTRL systems.

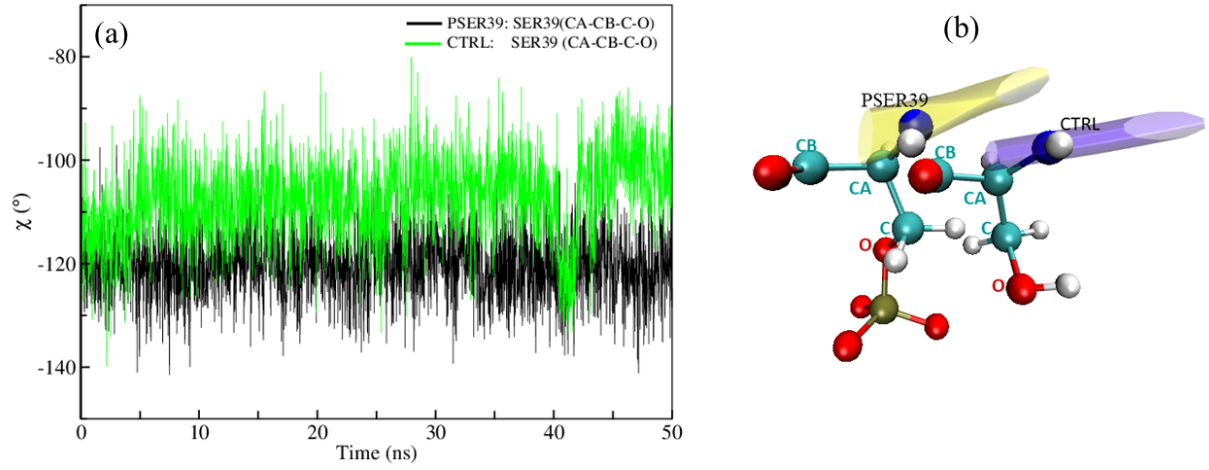


FIG. 2. (a) The variation of the $\chi$ dihedral angle of SER39 residue of Rap protein with time. The data is plotted for last 50ns of simulation data. (b) Representative snapshot of SER39 showing the dihedral angle ($\chi$ ) CB-CA-C-O corresponding to end structure of simulation for both system.

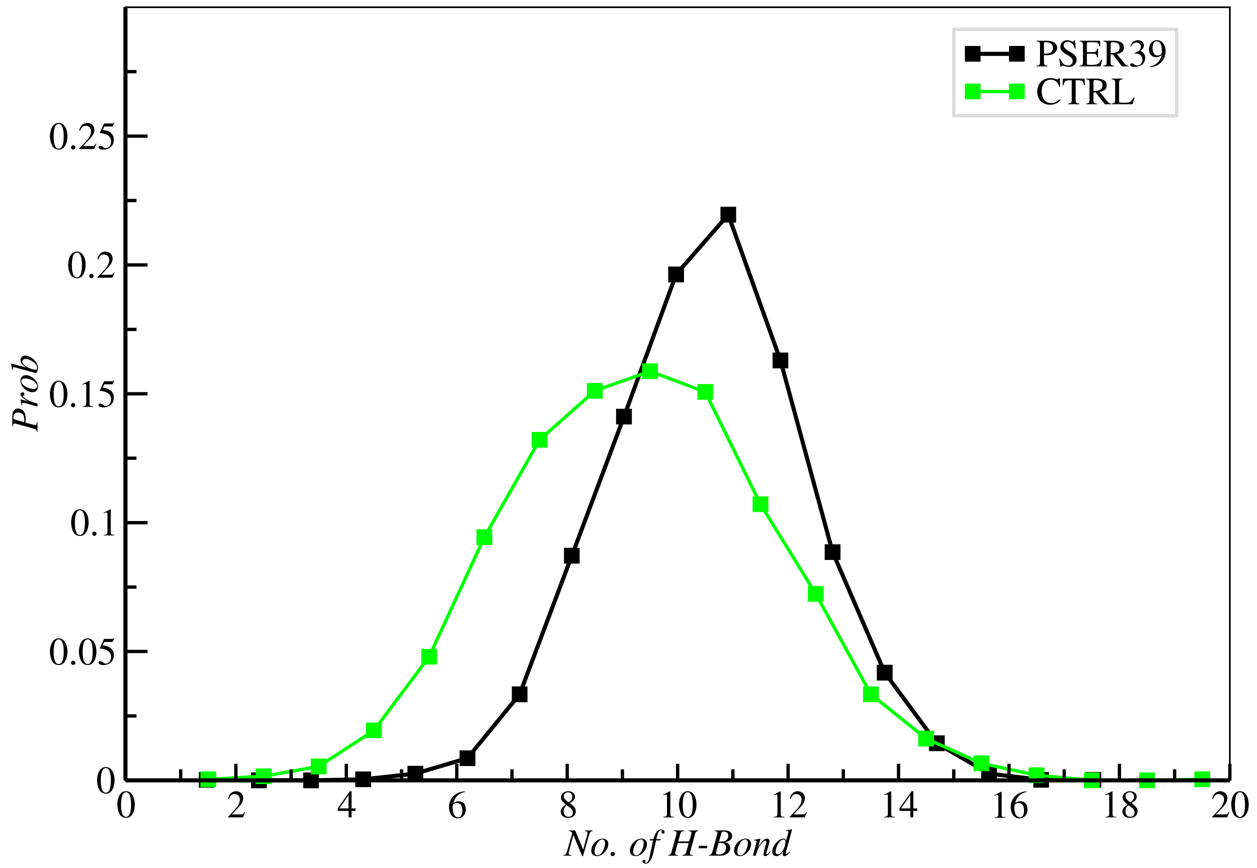


FIG. 3. The distribution of the no. of H-Bond formed between Rap and Raf protein calculated from last 100ns of simulation data for PSER39 and CTRL system.

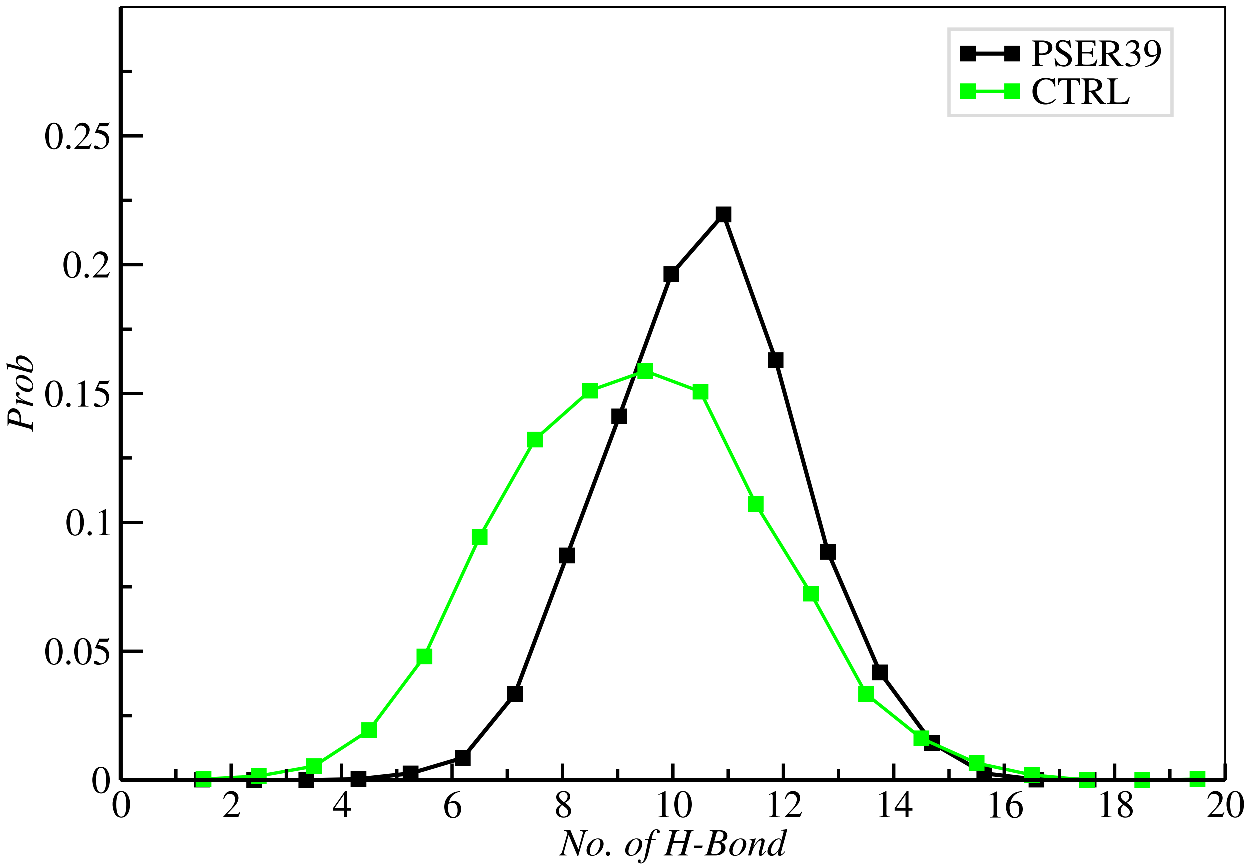


FIG. 4. The distribution of the no. of H-Bond formed between Rap and Raf protein calculated from last 100ns of simulation data for PSER39 and CTRL system.

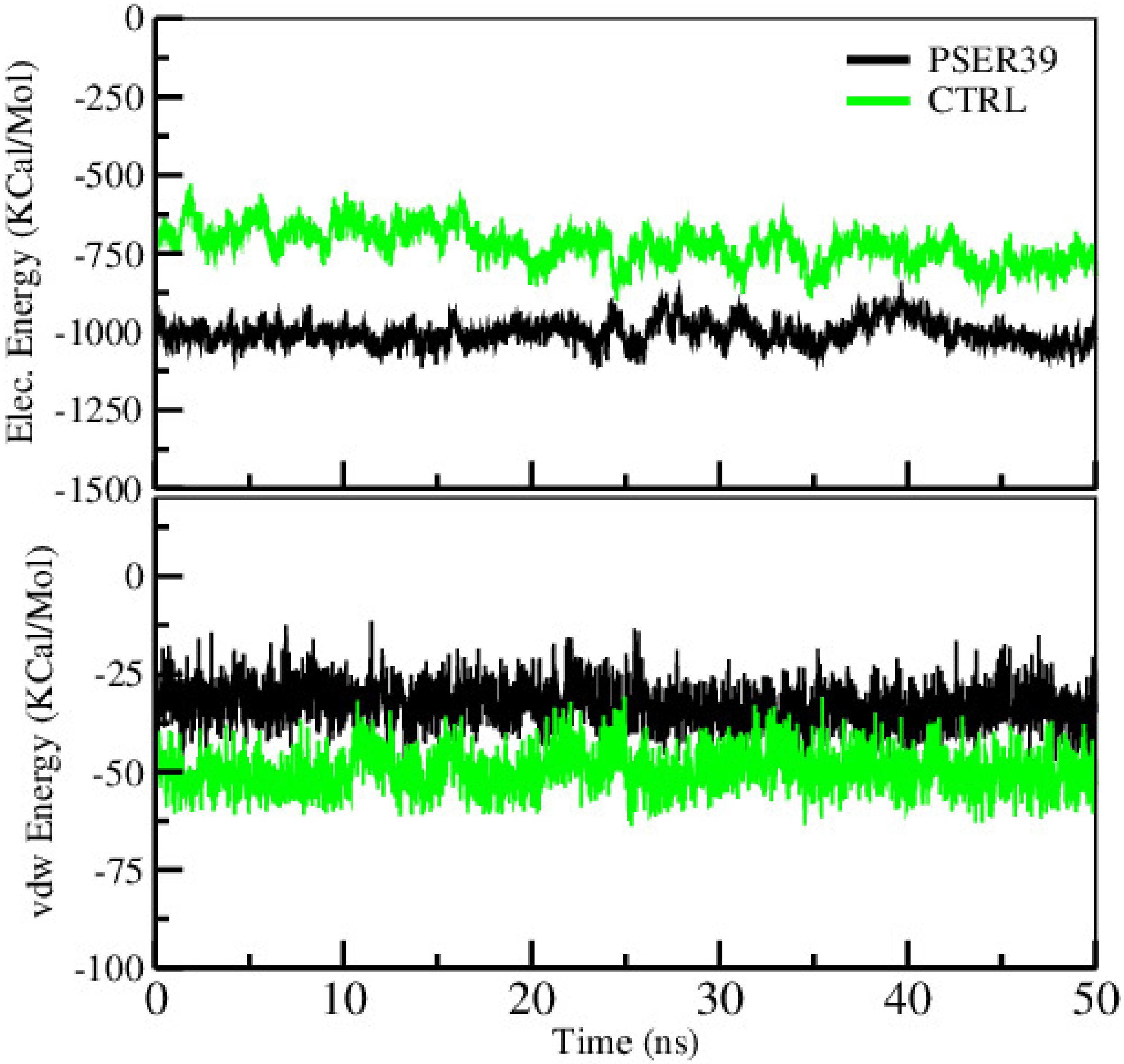


FIG. 5. Evolution of (a) electrostatic and (b) vdW interaction energies between Rap and Raf proteins in the last 50 ns of each simulation.
